# BNrich: A Bayesian network approach to the pathway enrichment analysis

**DOI:** 10.1101/2020.01.13.905448

**Authors:** Samaneh Maleknia, Ali Sharifi-Zarchi, Vahid Rezaei Tabar, Mohsen Namazi, Kaveh Kavousi

**Affiliations:** Laboratory of Complex Biological Systems and Bioinformatics (CBB),Department of Bioinformatics, Institute of Biochemistry and Biophysics, University of Tehran, Tehran, Iran; Department of Computer Engineering, Sharif University of Technology, Tehran, Iran; Department of Stem Cells and Developmental Biology, Cell Science Research Center, Royan Institute for Stem Cell Biology and Technology, Tehran, Iran; Department of Statistics, Allameh Tabataba’i University, Tehran, Iran; School of Biological Sciences, Institute for Research in Fundamental Sciences, Tehran, Tehran, Iran; Laboratory of Biomolecular Modeling (LBM), Department of Bioinformatics, Institute of Biochemistry and Biophysics, University of Tehran, Tehran, Iran

## Abstract

**Motivation:** One of the most popular techniques in biological studies for analyzing high throughput data is pathway enrichment analysis (PEA). Many researchers apply the existing methods without considering the topology of pathways or at least they have overlooked a significant part of the structure, which may reduce the accuracy and generalizability of the results. Developing a new approach while considering gene expression data and topological features like causal relations regarding edge directions will help the investigators to achieve more accurate results.

**Results:** We proposed a new pathway enrichment analysis based on Bayesian network (BNrich) as an approach in PEA. To this end, the cycles were eliminated in 187 KEGG human signaling pathways concerning intuitive biological rules and the Bayesian network structures were constructed. The constructed networks were simplified by the Least Absolute Shrinkage Selector Operator (LASSO), and their parameters were estimated using the gene expression data. We finally prioritize the impacted pathways by Fisher’s Exact Test on significant parameters. Our method integrates both edge and node related parameters to enrich modules in the affected signaling pathway network. In order to evaluate the proposed method, consistency, discrimination, false positive rate and empirical P-value criteria were calculated, and the results are compared to well-known enrichment methods such as signaling pathway impact analysis (SPIA), bi-level meta-analysis (BLMA) and topology-based pathway enrichment analysis (TPEA).

**Availability:** The R package is available on <u>carn</u>.

## 1 Introduction

Over the past few decades, the fast advances of high-throughput technologies have assisted the investigators to comprehensively monitor complicated biological systems. Subsequently, the innovative methods for all-inclusive surveys have been developed [1] providing a summary of the most advantageous information, all the while preserving and enhancing comprehensiveness. The pathway enrichment analysis (PEA) is one of the most popular among these methods [2–5].

The outcomes of the PEA methods can be lead to a definite and limited set of genes and intergenic relations through a pool of biological data, in order to perform later phases of research. These methods can be divided into three major categories under the chronologically sorted list of events [6]. The first generation is over-representation analysis (ORA) which uses just a subset of data (e.g., DEGs); inasmuch as the cut-off threshold is defined by the user, varying results might be obtained via ORA [7,8]. The second generation, the functional class scoring (FCS) methods, utilize all available expression data, even when information about the network structure of the pathways is missed [9–11]. The last generation, the topology-based pathway analysis (TPA), exploits biological (causal/non-causal) relation between pathway components [12–16]. In most cases, TPA methods provide a more accurate analysis [6]. However, a substantial proportion of these methods consider only the relationship between each gene and its neighbors [17–20] which ignores the complete topology of the network. Consequently, the methods are needed to deliberate the entire structure of the signaling pathways, as much as possible.

In this paper, we introduce BNrich, a new TPA method, based on the Bayesian Network (BN). The literature demonstrated that BN is a worthwhile technique for integrating and modeling biological data with causal relationships [21–25]. Actually, there are some pathway enrichment analysis methods using BN [26] or Dynamic BNs which are learned with time series gene expression data [27].

This study uses the BN to model variations in downstream components (children) as a consequence of change in upstream components (parents). The method employs signaling pathways as underlying structures for the inferred BNs [2,28,29] and gene expression data to estimate its parameters [26,30]. We eliminate cycles of inferred networks based on the biologically intuitive rules instead of using the computational algorithms [31]. In the BNrich, the edges that directly connect a gene product to itself or originated from the nucleus resulting in cycle formation, are removed. The inferred networks are simplified in two steps; unifying genes and LASSO. Likewise, we use originally continuous gene expression data rather than discretized data [26] to BN parameters learning. Given the methods of parameter learning in the BN, we estimate regression coefficients by continuous data [32,33]. The final impacted pathways are gained through Fisher’s exact test. This method can represent effective genes and biological relations in the impacted pathways based on the significance level.

We evaluate the proposed method on 187 human non-metabolic KEGG pathways using eight various gene expression datasets. Eventually, we compare the proposed framework with the other pathway topology-based methods, namely, SPIA [12], BLAM [15,34] and TPEA [16].

## 2 Methods

BNs as probabilistic graphical models that provide arguments in uncertainty framework by a directed acyclic graph (DAG) represent relations among random variables [32,35]. The nodes and arrows in BNs represent random variables and conditional dependence relations, respectively. Here, a group of experts manually proved these causal relationships in the labs [36]. The ability to represent prior data and knowledge in updatable relationships is one of the most important advantages of BNs; therefore, many researchers studied biological data benefiting BNs’ stated advantage [24,27,37].

Generally, reconstruction of BNs is implemented in two steps: structure learning and parameter learning. We first attained structures of BNs by KEGG pathways then simplified them. We estimated parameters using gene expression data via *maximum likelihood estimation* (MLE).

### 2.1 Reconstruction of BN Structures

The KEGG pathways are the collection of directed biological graphs whose biological causal relations have already been confirmed. Because they are not entirely acyclic, some of them require structural modification to eliminate the cycle. In this research these pathways become acyclic through a novel biological strategy. In the following, the simplification of the networks is performed with two steps of unifying genes and the LASSO regression to prepare the BN networks for estimating the parameters.

#### 2.1.1 Reconstruction of BNs’ structures utilizing signaling pathways

All the pathways of this study have been extracted from the KEGG pathways database^1^. They include all human, non-metabolic pathways in which gene products are as nodes and their numbers of edges vary between 10 to 1500 [19]. Totally, pathways included in this study are 187 of which 69 contain cyclic pathways. The elimination of the edges is accomplished through biological approach rather than computational methods [26]. The cyclic pathways have been converted to the acyclic graph while considering their biological perspective as follows:

- Edges that connect two nodes with the same label (gene product) were eliminated.
- According to the high importance of the cytoplasmic processes in this context, the feedbacks originating from nucleus resulting in cycle formation, were discarded.
- The main direction of signaling flux is from the cell membrane to the nucleus, in most of the cases. Therefore, in the remaining cycles, the edges which were opposite of the signaling flux direction and would lead to cycle configuration were removed.

The examples of these rules are displayed in supplementary (Figure S1). All 187 pathways were converted to a directed acyclic graph (DAG) after applying the above rules and eliminating 1,600 edges (less than 0.02 total edges); and we employed them as BNs structures.

#### 2.1.2 BN simplification

Though we extracted the structure of BNs directly from biological pathways, these structures need to be moderated and simplified before estimating the parameters. Thus, while maintaining the ability to estimate the correct parameters, inferencing from the final networks will also be more tractable [38]. This simplification takes place in two stages. The first step is unifying which utilizes classical BN pruning by deleting nodes for which gene expression data is not accessible depending on the dataset platform [39,40]. Similarly, genes that are not in the pathways are also removed from gene expression data to prepare the dataset for the parameter learning. Thereafter, the networks which have at least 5 edges remain in the analysis.

The second step of simplification is driven by the parameter learning model. In BN, the number of outgoing and incoming edges of a node shows the number of children and number of parents, respectively, and the expression level of the node is a function of expression levels of its parents [32].

The parameter learning method in BN depends on the distribution type of the variables including discrete, continuous and hybrid (Scutari, 2010). Here, continuous gene expression data is provided at two conditions of the case (disease or exposed) and control to learning BNs parameters. Hence, the mean value of gene expression for each node can be modeled as a linear regression of its parents’ gene expression. On the other hand, in many of the pathway databases including KEGG, a node is merely a position and can represent multiple gene products. Concurrently, many of these nodes are regarded as inputs of a single node such that some of them have up to 65 input nodes (parents). Given the type of estimation of parameters, this large number of parents means a large number of independent variables in the regression line and can lead to the problem of collinearity. To solve the problem, before the final estimation of the parameters, the LASSO regression is used to reduce the number of edges for each network [41,42]. LASSO as a variable selection and regularization method, minimizes the sum of squares of errors by an upper bound on the sum of absolute values of the model parameters as a solution to the l_1_ optimization problem [41,42]. This optimization problem can be written as an estimate of the parameter:

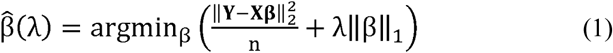

Where 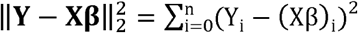 and 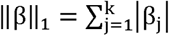 and λ ≥ 0 is the parameter that controls the strength of the penalty, the larger the value of λ, the greater amount of shrinkage [41]. The best value for λ was obtained with minimum mean error by using the cross validation (CV) procedure.

The LASSO method was applied for learning BN structure in some studies[43,44]. In this study, the Lasso regression is implemented on each of the remained networks from the previous stage for each dataset. We run LASSO regression for each of the aforesaid network structures for both case and control data. Afterwards, all edges which have zero coefficient for both networks are excluded. Due to computational concerns, the sample size should be greater than 20 across case and control samples.

### 2.2 BN Parameter learning

In the BN simplification phase, we use LASSO to resolve the collinearity problem using CV to obtain minimum mean error. The LASSO can be applied just on the regression lines with more than one independent variable, while at least a quarter of the nodes in signaling pathway networks are either roots or have just one parent, which means that they have at most one independent variable and hence LASSO cannot be applied to estimate their parameters. To have a unified parameter estimation technique for all nodes and noticing the fact that continuous gene expression data are exploited in the BNrich method [33,45] the maximum likelihood estimation (MLE) technique is carried out for parameter estimation. This estimation is done at two states of the case and control data which share identical structures. For each node *Y*, 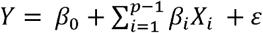, in which, X_1_ … X_p−1_ are the parents of *Y* [32]. In the network, *β*_1_ … *β*_*p*−1_, are interpreted as the weights of the edges and the amounts of biological functions based on the case and control data. The fixed value of *β*_0_ is the intercept of the regression line. In other words, the expected mean value of Y in case that all *X*_*i*_ (i = 1,2 …p − 1) are zero, is *β*_0_. Indeed, *β*_0_ is the amount of variable that is not modeled by its parents values.

### 2.3 Testing the equality of regression coefficients

Estimated Parameters of Bayesian Networks are derived from the following distribution [46]:

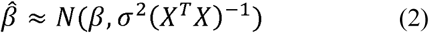

Note that this approximation can also be used for *β*_0_ by adding a column of ones to the matrix X. Statistic *σ* in above distribution is “standard deviation of the residuals” and was estimated by 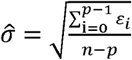, in which *n* is the number of samples and *p* is the number of parameters (*β*_*i*_), and *ε*_*i*_, is the i^th^ residual (*i* = 0,1,2 …*p* − 1)

Node Y has parents X_1_, X_2_… X_p−1_ in the network. When 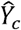 and 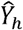 are the estimations of Y for the case and control datasets respectively which are modeled as follows:

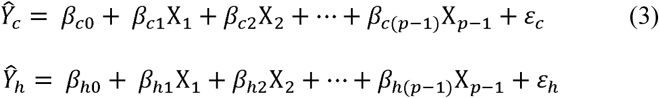

We compare corresponding *β* parameters between each pair of trained networks (for each pathway) in case and control states, by t-test [47] by degree of freedom as follows:

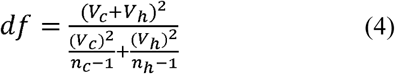

When 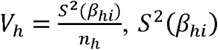, is estimated variance for *i*^*th*^ parameters of considering regression equation in control data, *n*_*h*_ is number of control individuals, 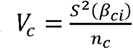, *S*^2^(*β* _*ci*_) is estimated variance for *i*^*th*^ parameters of considering regression equation in case (exposed or disease) data and *n*_*c*_ is number of case individuals. For the t-tests to yield comparable p-values, all variances and sample sizes are hypothesized to be different.

### 2.4 Prioritize the impacted pathways

We determine the enriched pathways by Fisher’s exact test which was utilized by many researchers for implementing enrichment analysis techniques [48–50]. In this stage, all paired parameters in case and control states are tested and significant parameters in any network are determined. Then, the Fisher’s exact test, as an overrepresentation test, is used for each network. The table of Fisher’s exact test for kth BN is as follows:

|  | #Significant<br>parameters | #Non significant<br>parameters |
| --- | --- | --- |
| #Parameters are in $k^{th}$ BN | $m_k$ | $n_k - m_k$ |
| #Parameters are NOT in $k^{th}$ BN | $M - m_k$ | $N - M - n_k + m_k$ |

*k* ∈ {1,2,3,…,*K*}; *K* is the number of remained networks,

*m*_*k*_= number of significant parameters in the *k*^*th*^ BN,

*n*_*k*_= number of all parameters in the *k*^*th*^ BN,

*M* = number of all significant parameters in all BNs,

*N* = number of all parameters in all BNs.

All of the trained BNs are examined by Fisher’s exact test, then significant pathways are determined as the results of the enrichment analysis. We calculate the adjusted P-value, by the Benjamini-Hochberg False Discovery Rate (FDR) correction method.

### 2.5 More insight into the impacted pathways

The impacted pathways can be visualized according to the significant threshold of the coefficients chosen by the user. The smaller threshold of the p-value (or FDR) results in the more compact pathway. The nodes’ and edges’ attributes in *k*^*th*^ pathway can be defined as below:

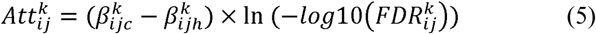

Where (*k* =1,…,K) in which K shows the number of final networks and 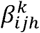 is the *j*^*th*^ parameter (coefficient) for *i*^*th*^ node in the healthy control data when *i* ∈ {1,2, …,*m*} and *j* ∈ {0,1,2, …,*p* − 1}. The parameter 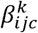 is the corresponding parameter (coefficient) of case data in the *k*^*th*^ pathway. 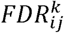 is also acquired from T-test between these two parameters.

As stated previously, the parameter *β*_*i*0(*h*/*c*)_ shows the *i*^*th*^ node attribute for (h)ealthy or (c)ase data. Subsequently, the attribute related to edge from node u to 1 is calculated by *β*_*ij*(*h*/*c*)_ when *j* ∈ {1,2, …*p* − 1}.

We exemplify a part of Platelet activation pathway enriched in the HT29 (Table1), the human colon cancer cell line dataset, to represent the advantage of the BNrich result visualization (section 3.3).

**Table 1.** Datasets for evaluation and comparison methods

| Dataset ID | Study | Tissue type | Sample size<br>(case/control) | Reference |
| --- | --- | --- | --- | --- |
| N/A | A375 | Tumor cell line skin | 110/110 | [55] |
| N/A | HT29 | Tumor cell line large intestine | 107/110 | [55] |
| GSE93601<sup>a</sup> | Breast Cancer | Tumor breast | 144/118 | [56] |
| GSE63060<sup>b</sup> | Alzheimer | Blood | 99/99 | [57] |
| GSE44076 | Colorectal Cancer | colon mucosa cells | 98/148 | [58] |
| GSE47756 | Colorectal Cancer | peripheral blood monocytes | 28/38 | [59] |
| GSE28000<sup>c</sup> | Colorectal Cancer | colon | 34/81 | [60] |
| GSE23878 | Colorectal Cancer | colorectal tissue | 35/24 | [61] |

a) Age at diagnosis $\geq 50$ ; estrogen receptor status: Positive; alcohol intake: 0 g/day
b) Age > 60; Alzheimer patients matched by healthy controls in age and sex
c) GPL4133

### 2.6 Evaluation

#### 2.6.1 Other methods for comparison

Three methods, i.e. TPEA, BLMA, and SPIA, were selected among the topology-based pathway analysis methods to be compared with the proposed method. The topology-based pathway enrichment analysis (TPEA) method, integrates topological properties and takes into account global upstream/downstream positions of genes in pathways [16]. BLMA [15,34] uses the gene expression data directly. It is worth mentioning that BLMA is able to analyze several datasets simultaneously. Finally, the signaling pathway impact analysis (SPIA) method combines the features of the classical enrichment analysis with measurements of the actual perturbation on a given pathway under a given condition [12];although this method is not new, it is still considered as a benchmark to examine new PEA methods [16,51,52].

#### 2.6.2 Evaluation criteria

We evaluate our method based on the four criteria (elaborated on below), namely, consistency, discrimination, False positive rate and empirical P-value by permutation test.

**\* Consistency** measures the number of overlapping nodes and edges among independent datasets under similar studying conditions, at a given rank threshold (top 10, 20… 70 significant pathways) [19,52]. We conduct consistency analysis on the last four datasets shown in Table1, related to colon cancer. This analysis has been performed in two phases: consistency analysis among four datasets and consistency analysis along dataset GSE44076 [52] by resampling 70% of data without replacement.
**\* Discrimination** measures the difference between results obtained from datasets originating from sufficiently different experimental conditions. These criteria are calculated by 100 times resampling without replacement for any subset 20%, 35%, 50%, 65%, 80% and 95% of datasets [53]. We perform discrimination analysis by first five datasets in Table1.
**\* False positive rate** is computed based on true target pathways in two cell line datasets. The cell line data is employed because their target pathways are more robust than the actual human data. Moreover, in contrast to the simulated data, the cell line data is biased towards neither the user preferences nor the evaluation method and leads to more generalized results.
**\* Empirical P-values** are attained using the permutation test on cell line datasets A375 (Table1) by generating 1000 simulated datasets in which we randomly assigned samples of the base dataset to two groups. This approach actually gives the observed type I error rate [46,54]. The empirical p-value is calculated by the formula below:

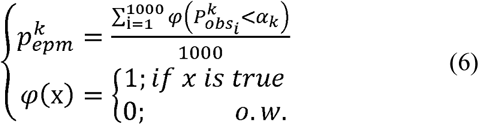

where 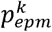 is empirical p-value based on *α*_*k*_, the real p-value of pathway *k* achieved by the BNrich., 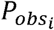 is the observed p-value obtained from the i^th^ permutation experiment for the k^th^ pathway and function *φ* counts the number of observed p-values which are less than *α*_*k*_ in permuted datasets.

## 3 Results

### 3.1 Dataset Preparation

We applied eight microarray gene expression datasets of 1383 samples from the NCBI database (Table 1). These datasets include two sets of cell line data [55]. We described characteristics of selected individuals and the target pathways related to the perturbation compounds, in supplementary (T1, Tabel S1 and Tabel S2).

### 3.2 BN structure simplification

As mentioned previously, simplifying BNs structures leads to a decrease in the number of edges and possibly networks. The number of edge reductions is different based on different dataset platforms according to the special set of gene expression afforded by each dataset platform. As it is shown in supplementary (Tabel S3), BNrich resulted in a different number of edges and networks in diverse dataset platforms.

### 3.3 Insight to the impacted pathways by visualization

As an instance of BNrich output, we illustrated a part of Platelet activation pathway enriched in HT29 cell line dataset visualized by Cytoscape 3.6.1(Figure 1).

**Fig1:**
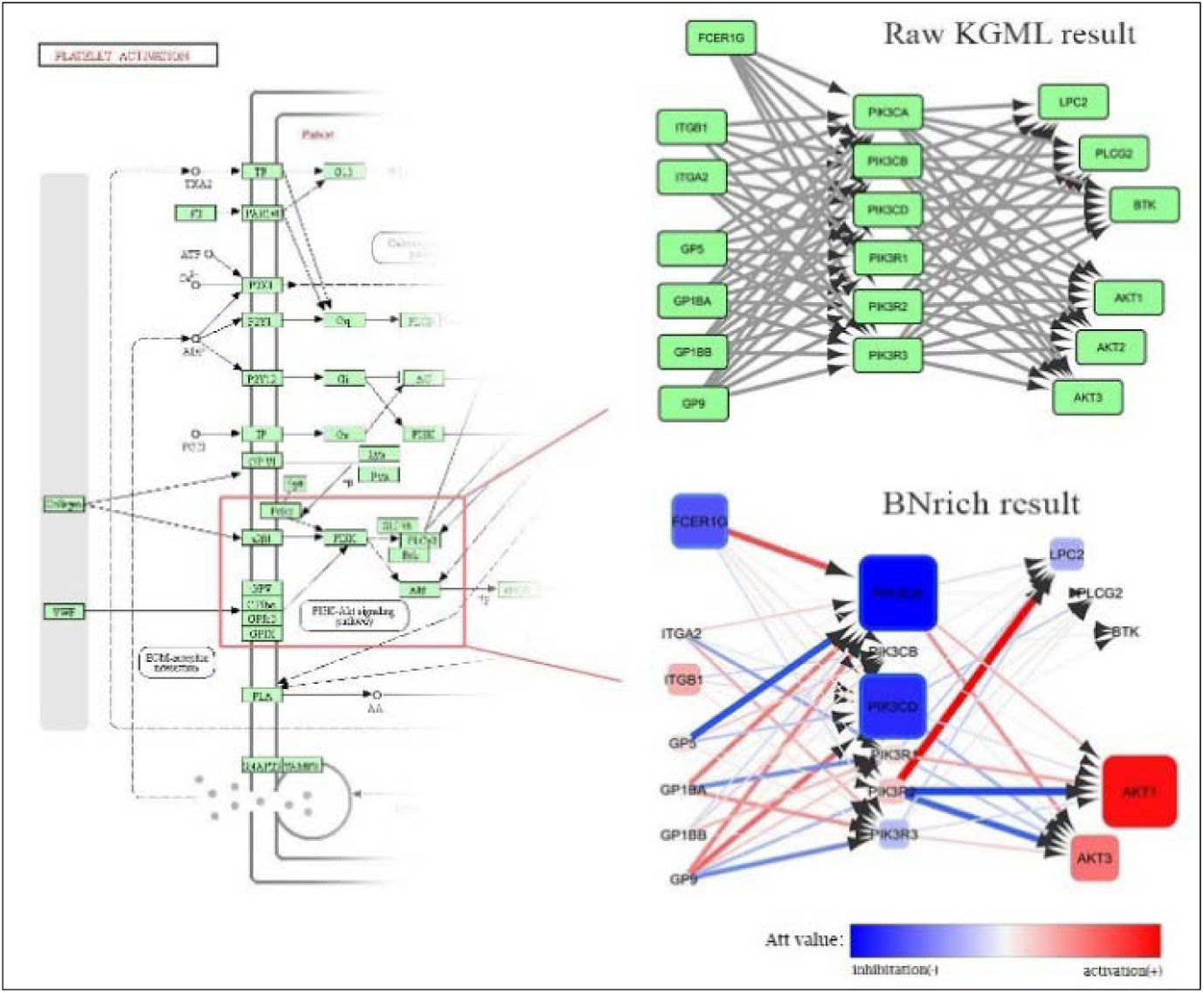
The Platelet activation pathway enriched in HT29 dataset. A part of the pathway related to PIK3 gene as a BNrich output is displayed. The color and size of nodes and edges reflect the values and the absolute values of the attribute (Att) respectively.

In accordance with BNrich method stages, at first, Platelet activation pathway was imported as a graphNEL graph with 282 directed edges and 124 nodes. After simplification, a BN with 219 directed edges and 109 nodes was attained. BNrich could determine 204 significant directed edges and 103 significant nodes.

### 3.4 Evaluation

#### 3.4.1 Consistency

We performed the methods of consistency on the four independent colorectal cancer datasets (the last four datasets in Table 1).

At the given number of top significant pathways, the results obtained by different methods on any pairwise datasets were shown in Figure 2. At the given FDR thresholds, the enriched pathways by different methods on any pairwise resampling datasets from GSE44076 were shown in supplementary (Figure S2).

**Fig2:**
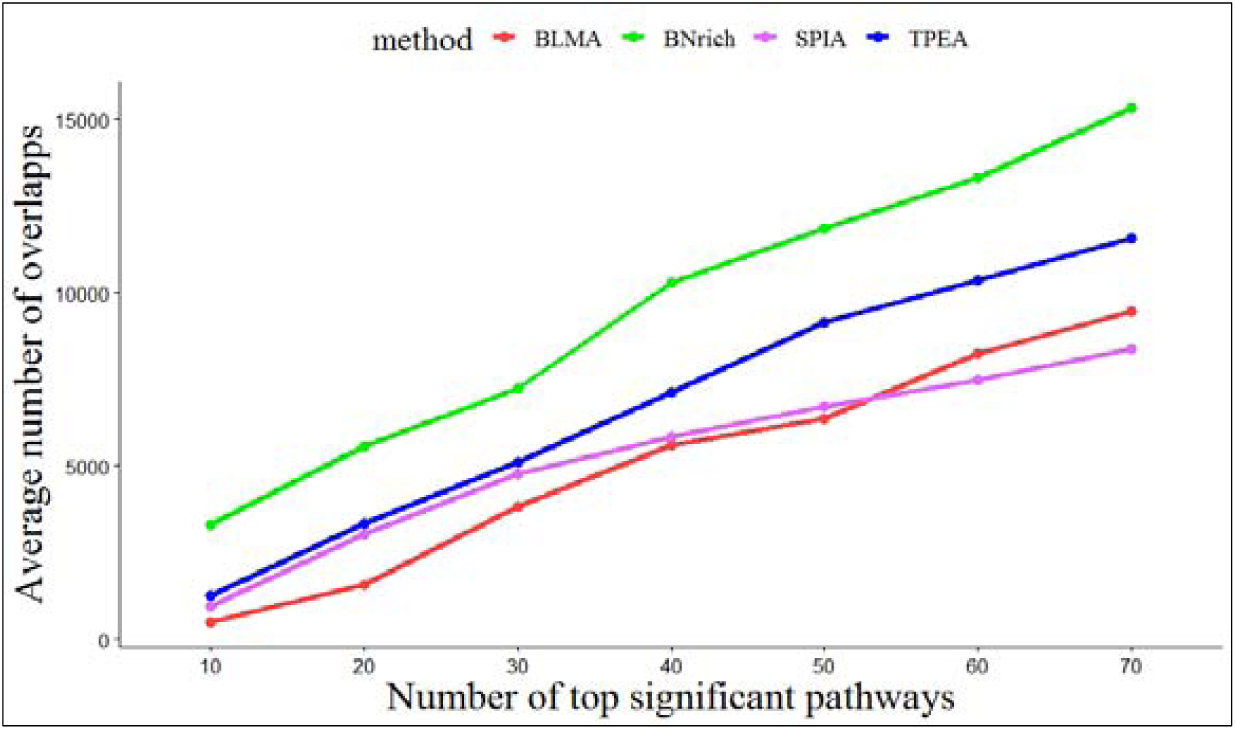
Plot of the average number of overlaps among impacted pathways based on the different number of top enriched pathways determined in four colorectal cancer datasets.

#### 3.4.2 Discriminant

The discriminant was computed by results of resampling datasets in four methods. Figure 3 shows the comparison of the discrimination values in GSE44076 vs HT29 datasets, among different methods. These results are achieved taking into account all the enriched pathways from different resampling levels of the first five datasets explained in Table1. All of the discriminant plots were shown in supplementary (Figure S3).

**Fig3:**
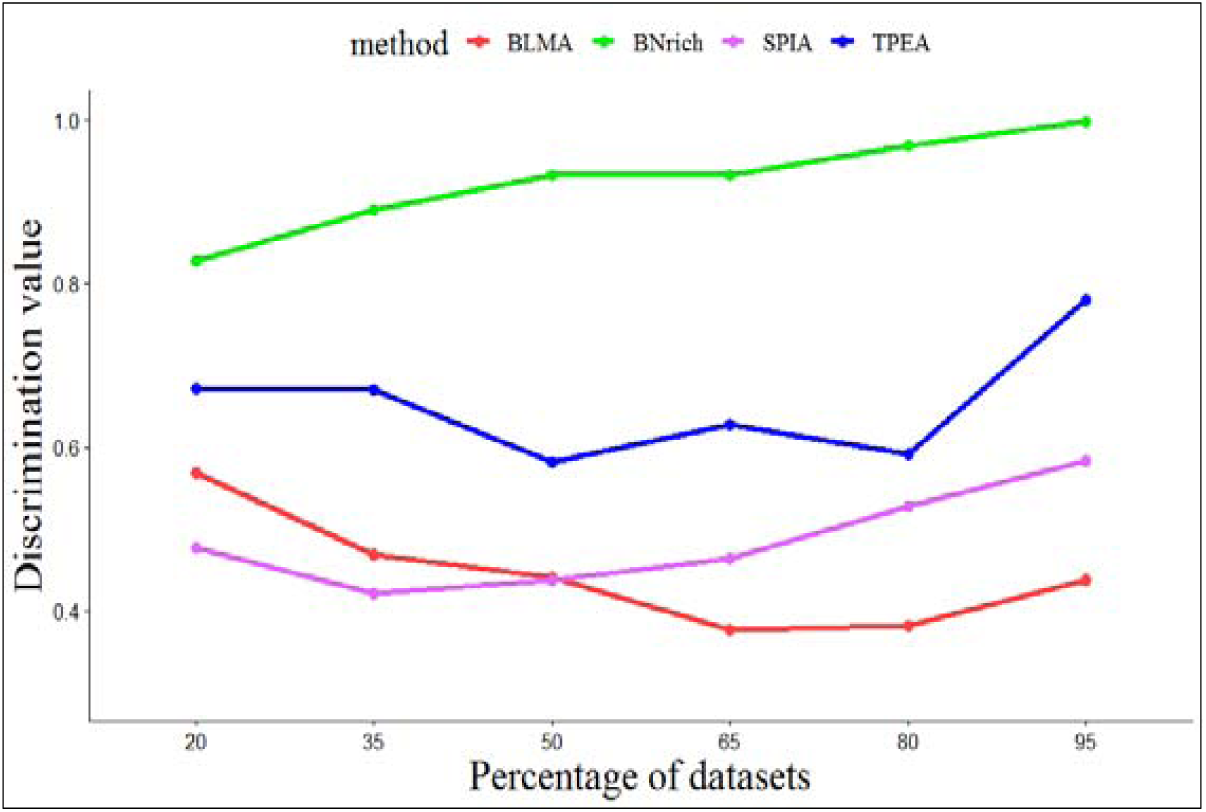
The discriminant values of various percentages of sampling among GSE44076 vs HT29

#### 3.4.3 False positive rate

As stated earlier, we calculated the false positive rate for two cell line datasets through determination of correct target pathways (Figure4).

**Fig4:**
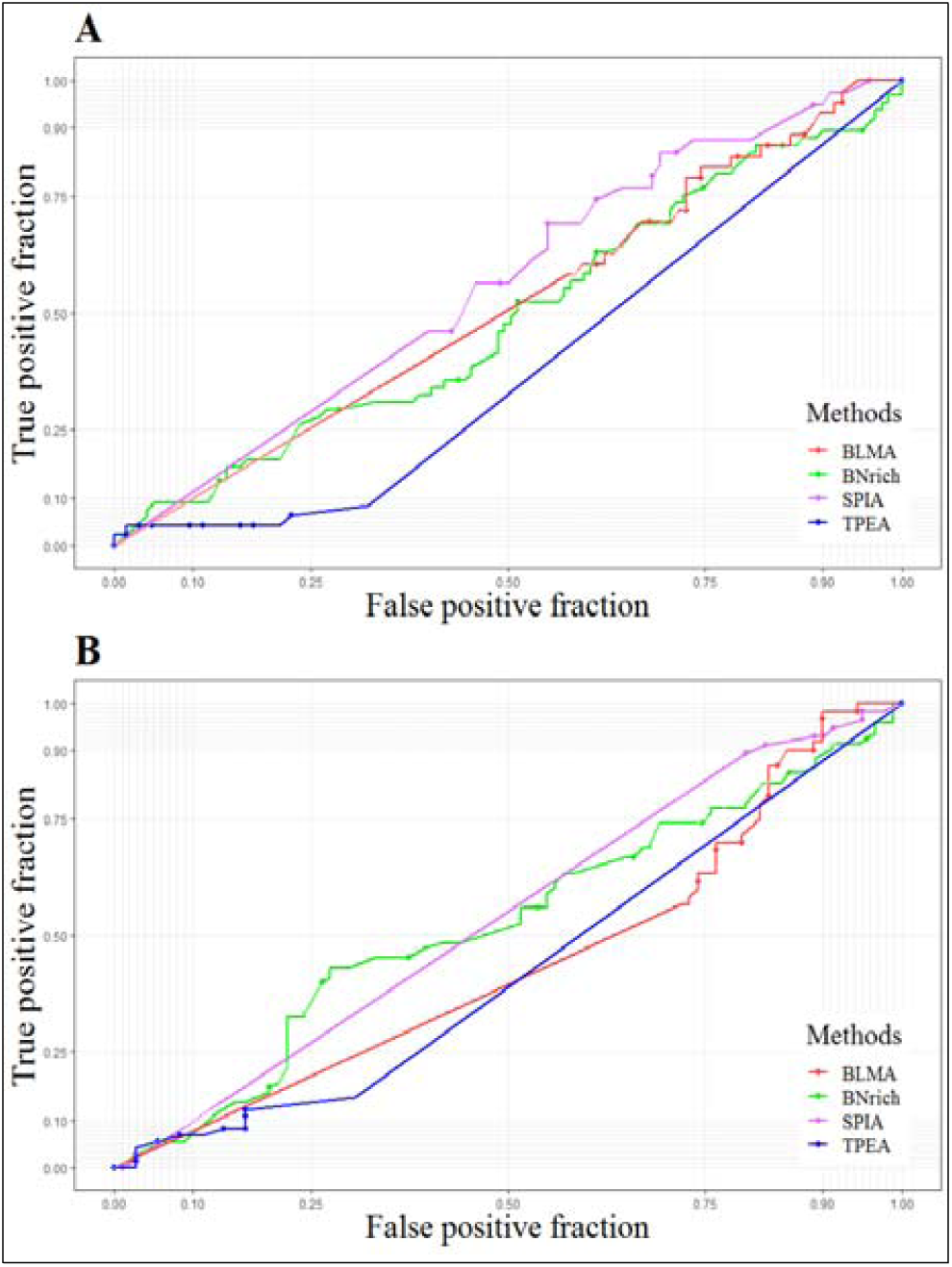
The ROC curves according to correct pathways in HT29 cell line dataset (A) and in A375 cell line dataset (B)

#### 3.4.4 Empirical P-value by permutation test

The empirical P-value was calculated for all pathways in dataset A375 is shown in Figure 5. According to formula 6, for ***α*** < **0.05**, the 75 percent of empirical P-values are less than 0.05 and are demonstrated by the red circle. Indeed, 47 percent of the dots are above the green line.

**Fig5:**
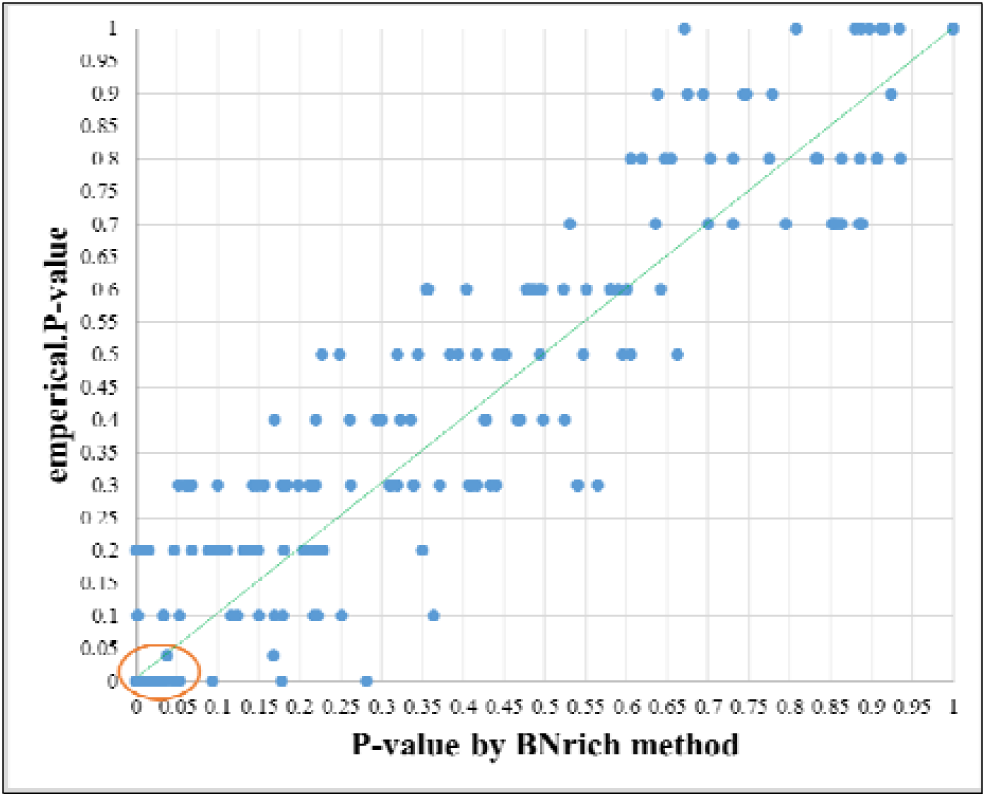
Empirical P-value vs. P-value by BNrich method. Among P-values less than 0.05 by BNrich method, 75% of empirical P-values are less than 0.05 and demonstrated by the red circle. Also, 47% of dots are the top of the green line.

## 4 Discussion

PEA enables investigators to benefit a mechanistic intuition into gene lists produced by genome-scale studies. These methods recognize biological pathways enriched in a gene list rather than anticipating them by chance [62]. On the other hand, the upstream regulators are generally causal genes in complicated diseases and control some downstream gene products; therefore, knowing these causal relationships can detect genes which are more likely to perform a key function in a pathway [16,63]. Thus, among the PEA methods, those that incorporate causal relationships into their computations are biologically preferred[6].

In the BNrich, the alteration of each component (gene product) is modeled by a linear regression equation with upstream components as regressors, in accordance with Bayesian network characteristics, while continuous gene expression data is applied to estimate parameters. Each gene expression variation is interpreted by two main factors: the level of changes in the gene’s parents and its own changes. The impact of changes which are related to *p* − 1 parents has been measured by *β*_1_…*β*_*p*−1_ parameters and illustrated as parameters of edges in BN. As previously stated, the presence of all edges (intergenic relations) has been already validated experimentally in vitro [36], and we only estimated their parameters in any gene expression dataset.

The effect of each individual component regardless of its parents has been shown by *β*_0_ parameter, which represents a node of BN. Consequently, both node and edge variations have been considered for the analysis, relying on the BN hierarchical framework. Further, we simplified the networks and excluded collinearity; consequently, we could decrease spatial/memory complexity without missing vital information via LASSO regression analysis.

The BNrich differs fundamentally from earlier methods. Among the earliest methods, one of the most famous ones is an enrichment analysis method using BNs [26], which may initially appear to be similar to our approach; however, there are, at least, two major differences.

The first difference is related to the modification of the BNs structure. Although in both methods, the BN structures are derived from signaling pathways, the afore-mentioned earlier method removed the cycles of these structures through a completely computational method introduced in 1995 [31] without any biological interpretation, while our method excludes some edges to eliminate cycles in the acquired networks through biological concepts.

Secondly, for learning BN structures, the BNrich, (i.e. our adopted method) utilizes continuous gene expression data, while it is the discretized fold-change data which is exploited in the competing earlier method. As a result, BNrich is more likely to potentially utilize transcriptome profiles. Certainly, another study is required to examine the extent to which the precision and accuracy of final results are affected by the information loss occurring during discretization.

The users can have a survey on enriched pathways in terms of both nodes and edges. For example, in Figure 1, we demonstrated a part of the Platelet activation pathway, which is enriched in HT29 cell line dataset. This cell line was exposed to perturbation compound, Dasatinib, an outcome of which is inhibiting PIK3. PIK3 gene has six paralogs which have been affected differentially by the compound. The colors of the nodes and edges are related to the values of the attribute (Att), while their size is related to absolute values of Att. Thence, among these paralogs, PIK3CA and PIK3CD were the most inhibited paralogs, while PIK3R2 had the strongest biological relations with the next genes in this pathway. It is worth mentioning that these effects are just related to this pathway. Likewise, the researchers need to examine the genes and their relationships in all the enriched pathways in order to gain more accurate insights.

The method of interest was evaluated through a variety of approaches. Figure 2 demonstrated that BNrich is more consistent compared to other methods, and it can identify similar pathways from same clinical condition data in different platforms of datasets. In Figure S2, the BNrich, after TPEA method, is also more consistent than the other two methods. Of course, it should be noted that the number of overlapping nodes and edges is directly related to the number of impacted pathways; as a result, inasmuch as the TPEA method produces many false positives in each run especially in a large dataset, this output is not unexpected.

Considering the second criteria, namely discrimination, BNrich method could better discriminate among specific experimental states in diverse studies; furthermore, with reference to false positive rate, our method results were quite comparable with those of previous reliable studies.

The results of the permutation test revealed that the accuracy of P-value in the introduced method is satisfactory. The mean of empirical P-value in each level (0.01, 0.05, 0.1 …) is at least equal to the P-value of the method. The equality of these two values means that BNrich performs as accurately as an exact test. Knowing that the last stage of the method was conducted using Fisher’s exact test, this conclusion was expected. Generally, accuracy of BNrich is not be any means less than traditional and acceptable tests.

### 4.1 Concluding Remarks

The main advantages of the BNrich, as a biologically intuitive method, include: exploiting the structural information acquired from signaling pathways, such as causal relationships between genes, to achieve more precise pathway enrichment in comparison with current well-known methods; controlling the false positive rate in networks with the LASSO regression [64]; applying a bi-stage simplification method for inferring the final structure of networks; and Using an integrative approach which considers both edge and node related parameters to enrich modules in the affected signaling networks.

### 4.2 Suggestions for Further Research

One way to upgrade the desired algorithm is applying additional information related to phenotype, with the purpose of assigning weigh to edges of the signaling pathways. It can lead to an increase in the speed of execution and a rise in the accuracy of the results while implementing LASSO regression.

## Supporting information

Fig S2: Plot of the average number of overlaps among impacted pathways determined in resampled datasets of a colorectal cancer dataset

Fig S3: The discriminant values of various percentages of sampling

ig S1: Some of the edges in the imported pathways eliminated by the biological approach

## Acknowledgements

The authors would like to thank Dr. Marco Scutari from Institute Dalle Molle of Studies on Arti-ficial Intelligence (IDSIA) and Mao Tanabe from Institute of Chemical Research, Kyoto Univer-sity, for their expert insights and suggestions shared via several emails contributing greatly to the quality of this research.

## Footnotes

1 KEGG PATHWAY Database available in: https://www.kegg.jp/kegg/pathway.html (Release 90.0, April 1, 2019)

