## Supplementary material for "BNrich: A Bayesian network approach to the pathway enrichment analysis": Fig S2: Plot of the average number of overlaps among impacted pathways determined in resampled datasets of a colorectal cancer dataset

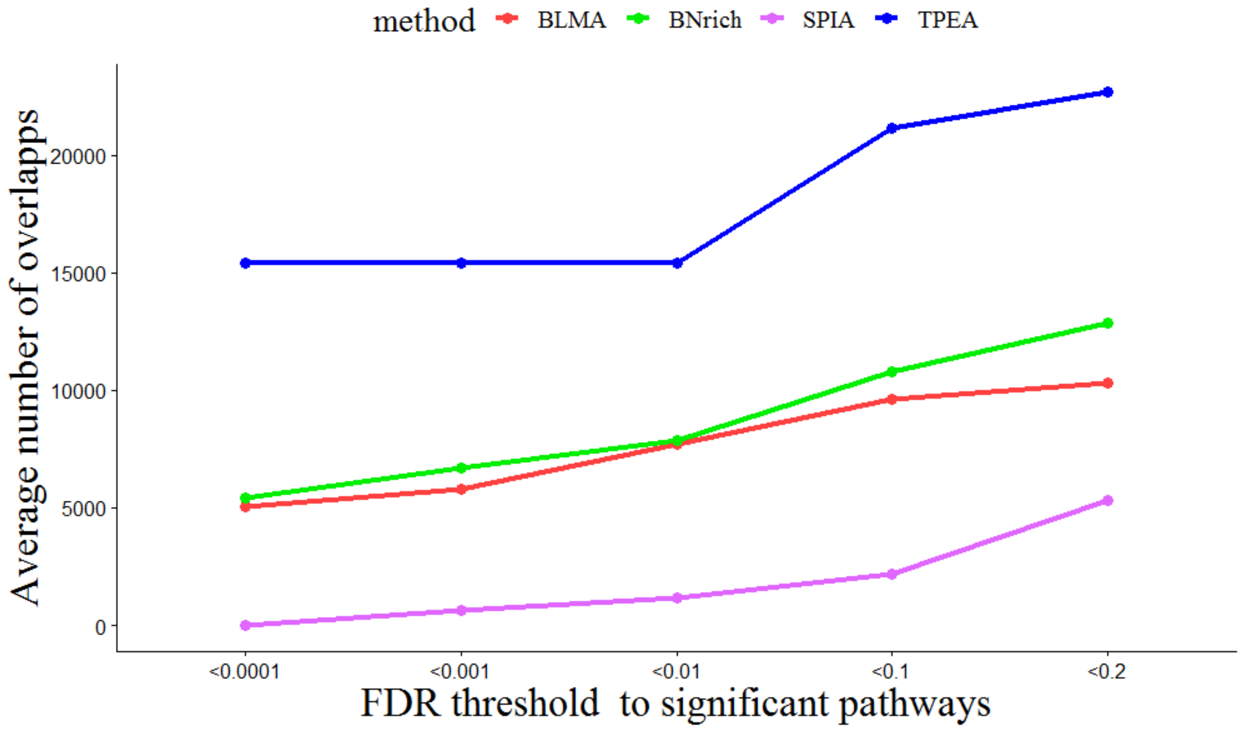


Fig S2: Plot of the average number of overlaps among impacted pathways determined in resampled datasets of a colorectal cancer dataset (GSE44076) at the different threshold of FDR
