## Supplementary material for "BNrich: A Bayesian network approach to the pathway enrichment analysis": Fig S3: The discriminant values of various percentages of sampling

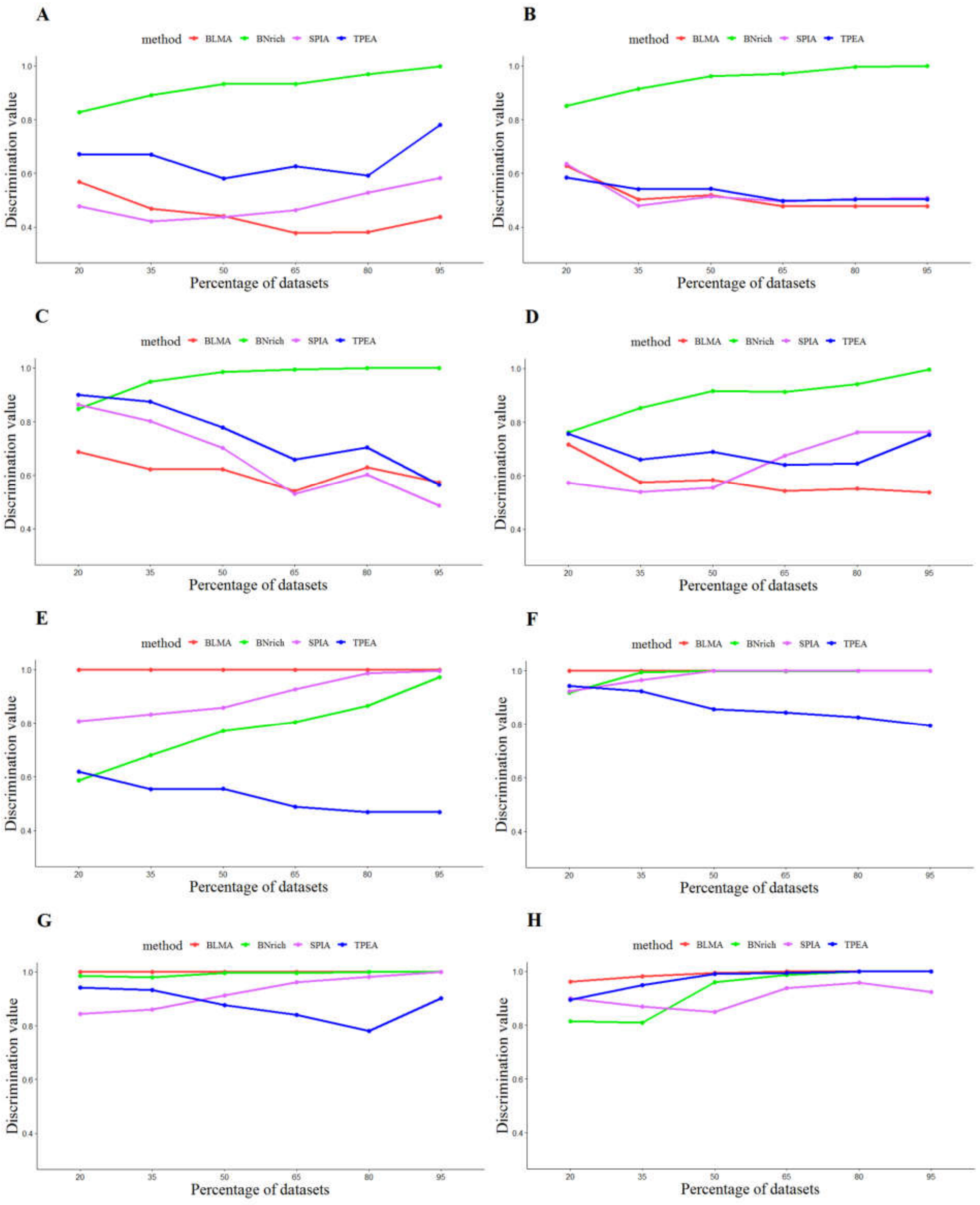


Fig S3: The discriminant values of various percentages of sampling among GSE44076 vs HT29 (A), GSE93601 vs GSE44076 (B), GSE44076 vs GSE63060(C)A375 vs GSE44076 (D), A375 vs GSE93601(E), HT29 vs GSE63060 (F), GSE63060 vs A375(G) and A375 vs HT29 (H)
