## Supplementary material for "BNrich: A Bayesian network approach to the pathway enrichment analysis": ig S1: Some of the edges in the imported pathways eliminated by the biological approach

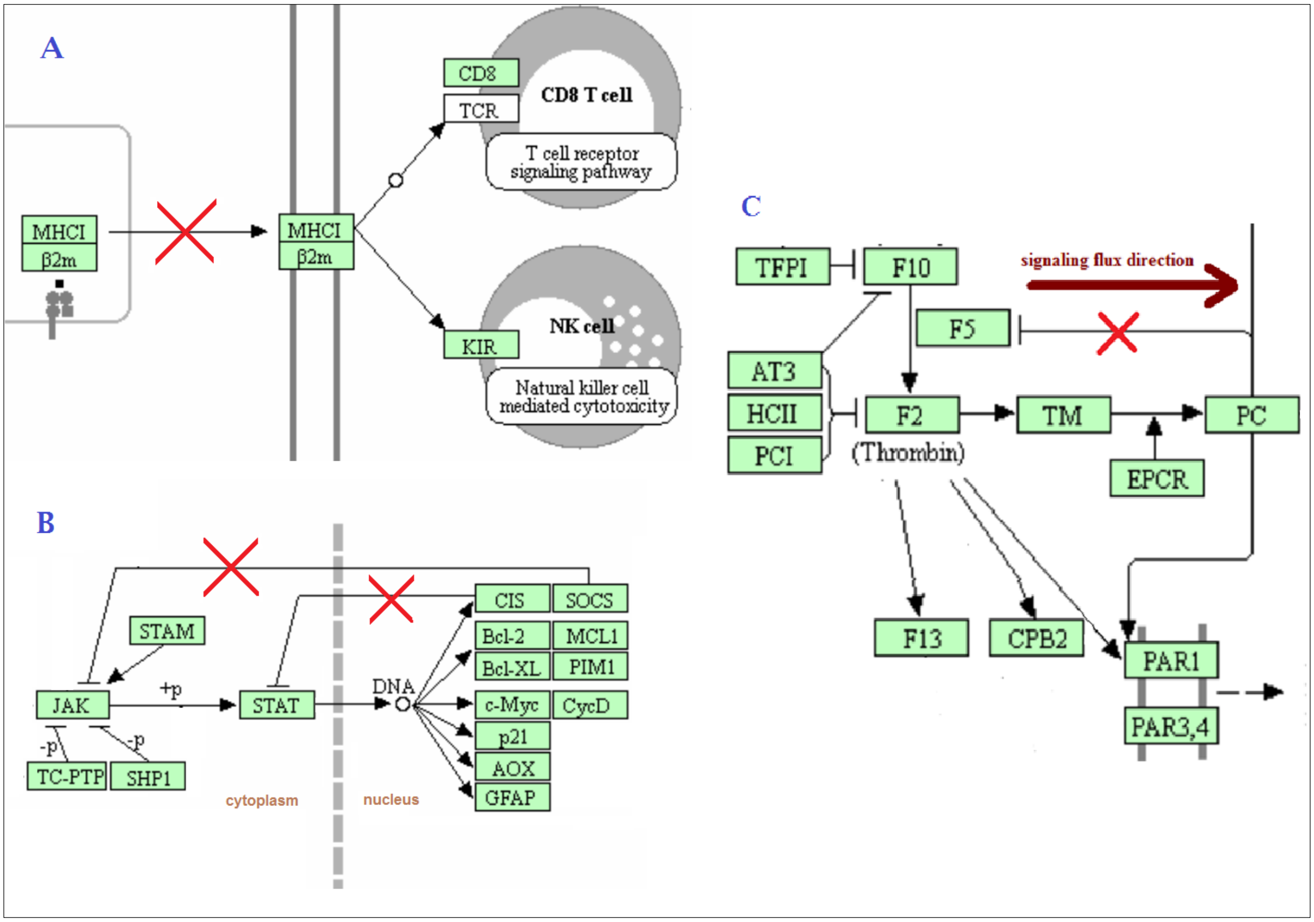


Fig S1: Some of the edges in the imported pathways eliminated by the biological approach, for example in the Antigen processing and presentation pathway, the edge connecting MHCI node to itself (A) and two edges in JAK-STAT signaling pathway regard-ing to the second declared rule (B) were removed. In the complement and coagulation cascades, the edge is the opposite of signaling flux direction, is excluded (C).
