## Supplemental Data 1 for "BNrich: A Bayesian network approach to the pathway enrichment analysis"

**T1. The characteristics of individuals and target pathways in cell line datasets**

We selected these two datasets in *cell_id* are A375 and HT29 by the following characteristics in *GSE70138_Broad_LINCS_cell_info* file:

1. Original cell line: it means *base_cell_id* = cell_id
2. *original_growth_pattern* was adherent
3. Sampling tissue was not missing: it means *primary_site* ≠ -666
4. Control individuals were selected randomly of those *pert_iname* = DMSO. Their number in each dataset is proportional to the number of case (exposed) individuals.

The perturbation compounds are PD-0325901 and Dasatinib respectively, those are FDA approved and also their functions were identified by *PubChem* or *KEGG drug* database. Among these filtered data, we selected cell lines line specificity > 0, cell_tas > 0.75.

We achieved target genes or modules for any perturbagen by The CLUE API playground. Afterwards, the target pathways were determined by the following techniques:

1- Searching *pert_iname* in *KEGG drug* database and find *Target Pathway*
2- Searching target genes or modules in *KEGG NETWORK* to find target pathways
3- Searching target genes in KEGG GENE database to find target pathways

The target pathways are as fallow:

**Table S1: The target pathways in A375 datasets in alphabetical order**

| | | |
|---|---|---|
| Acute myeloid leukemia | Focal adhesion | Osteoclast differentiation |
| Adherens junction | FoxO signaling pathway | Oxytocin signaling pathway |
| Alcoholism | Gap junction | Pancreatic cancer |
| Amyotrophic lateral sclerosis (ALS) | Gastric cancer | Pathways in cancer |
| Apelin signaling pathway | Glioma | Phospholipase D signaling pathway |
| Apoptosis | GnRH signaling pathway | PI3K-Akt signaling pathway |
| B cell receptor signaling pathway | Hepatitis B | Platinum drug resistance |
| Bladder cancer | Hepatitis C | Prion diseases |
| Breast cancer | Hepatocellular carcinoma | Progesterone-mediated oocyte maturation |
| Calcium signaling pathway | Herpes simplex infection | Prolactin signaling pathway |
| cAMP signaling pathway | HIF-1 signaling pathway | Prostate cancer |
| Cellular senescence | Human papillomavirus infection | Proteoglycans in cancer |
| Central carbon metabolism in cancer | Human T-cell leukemia virus 1 infection | Rap1 signaling pathway |
| cGMP-PKG signaling pathway | IL-17 signaling pathway | Ras signaling pathway |
| Chagas disease (American trypanosomiasis) | Inflammatory mediator regulation of TRP | Regulation of actin cytoskeleton |
| Chemokine signaling pathway | Influenza A | Relaxin signaling pathway |
| Choline metabolism in cancer | Insulin signaling pathway | Renal cell carcinoma |
| Cholinergic synapse | Intestinal immune network for IgA | RIG-I-like receptor signaling pathway |
| Chronic myeloid leukemia | Kaposi sarcoma-associated herpesvirus | Serotonergic synapse |
| Colorectal cancer | Leishmaniasis | Signaling pathways regulating pluripotency of |
| C-type lectin receptor signaling pathway | Long-term depression | Sphingolipid signaling pathway |
| Cushing syndrome | Long-term potentiation | T cell receptor signaling pathway |
| EGFR tyrosine kinase inhibitor resistance | MAPK signaling pathway | Thermogenesis |
| Endocrine resistance | Measles | Thyroid cancer |
| Endometrial cancer | Melanogenesis | Thyroid hormone signaling pathway |
| Epithelial cell signaling in Helicobacter pylori infection | Melanoma | TNF signaling pathway |
| Epstein-Barr virus infection | mTOR signaling pathway | Toll-like receptor signaling pathway |
| ErbB signaling pathway | Natural killer cell mediated cytotoxicity | Toxoplasmosis |
| Estrogen signaling pathway | Neurotrophin signaling pathway | Vascular smooth muscle contraction |
| Fc epsilon RI signaling pathway | NOD-like receptor signaling pathway | VEGF signaling pathway |
| Fc gamma R-mediated phagocytosis | Non-alcoholic fatty liver disease (NAFLD) | Wnt signaling pathway |
| Fluid shear stress and atherosclerosis | Non-small cell lung cancer | |

**Table S2: The target pathways in HT29 datasets in alphabetical order**

| | | |
|---|---|---|
| Acute myeloid leukemia | Focal adhesion | Pathogenic Escherichia coli infection |
| Adherens junction | GABAergic synapse | Pathways in cancer |
| Axon guidance | Gap junction | Phospholipase D signaling pathway |
| B cell receptor signaling pathway | Glioma | PI3K-Akt signaling pathway |
| Bacterial invasion of epithelial cells | GnRH signaling pathway | Platelet activation |
| Bladder cancer | Hematopoietic cell lineage | Primary immunodeficiency |
| Breast cancer | Hepatitis B | Prion diseases |
| Calcium signaling pathway | Herpes simplex virus 1 infection | Prolactin signaling pathway |
| Cell cycle | Human cytomegalovirus infection | Prostate cancer |
| Central carbon metabolism in cancer | Human papillomavirus infection | Proteoglycans in cancer |
| Chemokine signaling pathway | Human T-cell leukemia virus 1 infection | Rap1 signaling pathway |
| Choline metabolism in cancer | Inflammatory mediator regulation of TRP channels | Ras signaling pathway |
| Cholinergic synapse | Jak-STAT signaling pathway | Regulation of actin cytoskeleton |
| Chronic myeloid leukemia | Kaposi sarcoma-associated herpesvirus infection | Relaxin signaling pathway |
| C-type lectin receptor signaling pathway | Long-term depression | Shigellosis |
| EGFR tyrosine kinase inhibitor resistance | MAPK signaling pathway | Sphingolipid signaling pathway |
| Endocrine resistance | Melanogenesis | T cell receptor signaling pathway |
| Endocytosis | Melanoma | Th1 and Th2 cell differentiation |
| Epithelial cell signaling in Helicobacter pylori infection | MicroRNAs in cancer | Th17 cell differentiation |
| Epstein-Barr virus infection | Mitophagy - animal | Thyroid hormone signaling pathway |
| ErbB signaling pathway | Natural killer cell mediated cytotoxicity | Tight junction |
| Estrogen signaling pathway | Neurotrophin signaling pathway | Tuberculosis |
| Fc epsilon RI signaling pathway | NF-kappa B signaling pathway | VEGF signaling pathway |
| Fc gamma R-mediated phagocytosis | Osteoclast differentiation | Viral carcinogenesis |
| Fluid shear stress and atherosclerosis | Oxytocin signaling pathway | Viral myocarditis |
